# A Long-Term *Drosophila* Body Mass Measurement System

**DOI:** 10.1101/2025.05.29.656772

**Authors:** Yangyuan Li, Dan Feng, Wanhe Li

## Abstract

We propose a novel system for long-term monitoring of *Drosophila* body mass. In this approach, a cantilever beam is placed inside a *Drosophila* culture vial and undergoes random vibrations induced by *Drosophila* landings. An infrared camera connected to a microcomputer records these vibrations. Image processing techniques then extract the beam’s vibration signals from the video recordings. Applying the Euler-Bernoulli beam theory, we calculate the *Drosophila* body mass. As a demonstration, we used this system to measure body mass variations in wild-type *Drosophila* over 14 days.

**Summary statement:** This study presents a system that enables long-term monitoring of fruit fly body mass, offering new opportunities to study relationships between genes, circadian rhythms, phenotype, and long-term mass variation.

## INTRODUCTION

*Drosophila melanogaster* is a widely used model organism in biological, genetic, and medical research due to its short life cycle, ease of genetic manipulation, and physiological similarities to humans. Numerous studies have investigated *Drosophila*’s behaviors and physiological phenotypes using various methods. Widely employed techniques include infrared sensors for tracking circadian activity (e.g., Drosophila Activity Monitor, DAM, Trikenetics Inc.); cameras for movement and flight behavior analysis (Fontaine et al., 2009; Zimmerman et al., 2008; Gilestro, 2012); and artificial intelligence combined with imaging techniques for detailed behavior identification (Günel et al., 2019; Mathis et al., 2018; Pereira et al., 2019). However, despite body weight being an important metric for assessing physiological status (Harvey and Pagel, 1991; Peters, 1986; Reiss, 1989), a reliable and cost-effective method for long-term *Drosophila* body mass measurement is currently lacking. This limitation arises primarily from the challenge of measuring the very small *Drosophila* body mass (typically 0.1 mg to 1 mg; Burggren et al., 2017; Jumbo-Lucioni et al., 2010), as well as the high cost of existing equipment. Conventional body mass measurement relies on electronic balances. While accurate, they are unsuitable for long-term monitoring due to their high cost, susceptibility to environmental disturbances, and the need to repeatedly anesthetize and transfer individual flies—a process that can significantly affect their physiological state (Shen et al., 2019). Although alternative methods like laser displacement sensors (Shimazaki et al., 2022) and MEMS force sensors (Bartsch et al., 2007; Kohyama et al., 2018; Takahashi et al., 2014) have demonstrated potential for measuring *Drosophila* body mass, they also fall short for long-term body measurement monitoring of individual animals due to their high costs, susceptibility to environmental disturbances, and inability to measure over extended periods.

Here, we present a novel system for long-term *Drosophila* body mass monitoring that integrates a microcomputer, an infrared camera, image processing, and cantilever beams. This system involves placing a cantilever beam inside a *Drosophila* culture vial and using a Raspberry Pi 5 with an infrared camera to capture the vibrations of the cantilever beam induced by the fly’s movements. Image processing techniques, including edge detection and background subtraction, are then employed to extract the beams’ vibration data and the fly’s positional information. The body mass of the fly is subsequently calculated based on the vibrations of the cantilever beam. This system is cost-effective, with a single monitoring unit capable of tracking six flies simultaneously at an approximate cost of $70. It also provides a stable living environment that does not disrupt the flies’ natural behavior. We validated the system’s stability through continuous mass monitoring experiments involving multiple flies over several weeks.

## MATERIALS AND METHODS

### Fly husbandry and experimental setup

The wild-type Canton-S (CS) strain of *Drosophila melanogaster* was used in this study. Flies were raised on standard cornmeal-molasses-yeast-agar food at room temperature. Six individual flies were each placed in separate vials containing fly food and the cantilever beam (Fig. S1A), which were then mounted on the monitoring device (Fig. S1A) for high-frame-rate video recording. The system utilizes a Raspberry Pi 5 to stably capture video at 1280×720 pixels resolution and 120 frames per second (fps).

### Vibration signal extraction

Image processing techniques were used to extract the beam’s vibration frequencies and the fly’s positions. We first employed edge detection and connected component labeling to isolate the beam’s motion signals. To enhance processing efficiency, we first defined regions of interest, as indicated by rectangular boxes in Fig. 1A. Within each region of interest, we applied the Canny edge detector (Canny, 1986) and connected component labeling (‘bwlabel,’ MATLAB, Version R2024b, The MathWorks, Natick, MA, USA). From the detected connected components, we selected those with a large span in the *x*-direction to represent the beam’s edge (Fig. 1B). Vibration signals were extracted from the region corresponding to 30% of the cantilever beam’s length. This region was divided into 8 sections, and the centroid coordinates of each section were computed (marked with red Xs in Fig. 1C). Subsequently, based on the fly’s position (extracted in a later step), centroid coordinates were selected according to the following criteria: (1) if the fly did not obstruct the beam (Case 1, Fig. 1D), all points were selected; (2) if the fly partially obstructed the beam (Case 2, Fig. 1D), the points closest to the fly were unselected. Only the selected points were used for vibration signal extraction.

**Figure 1.**
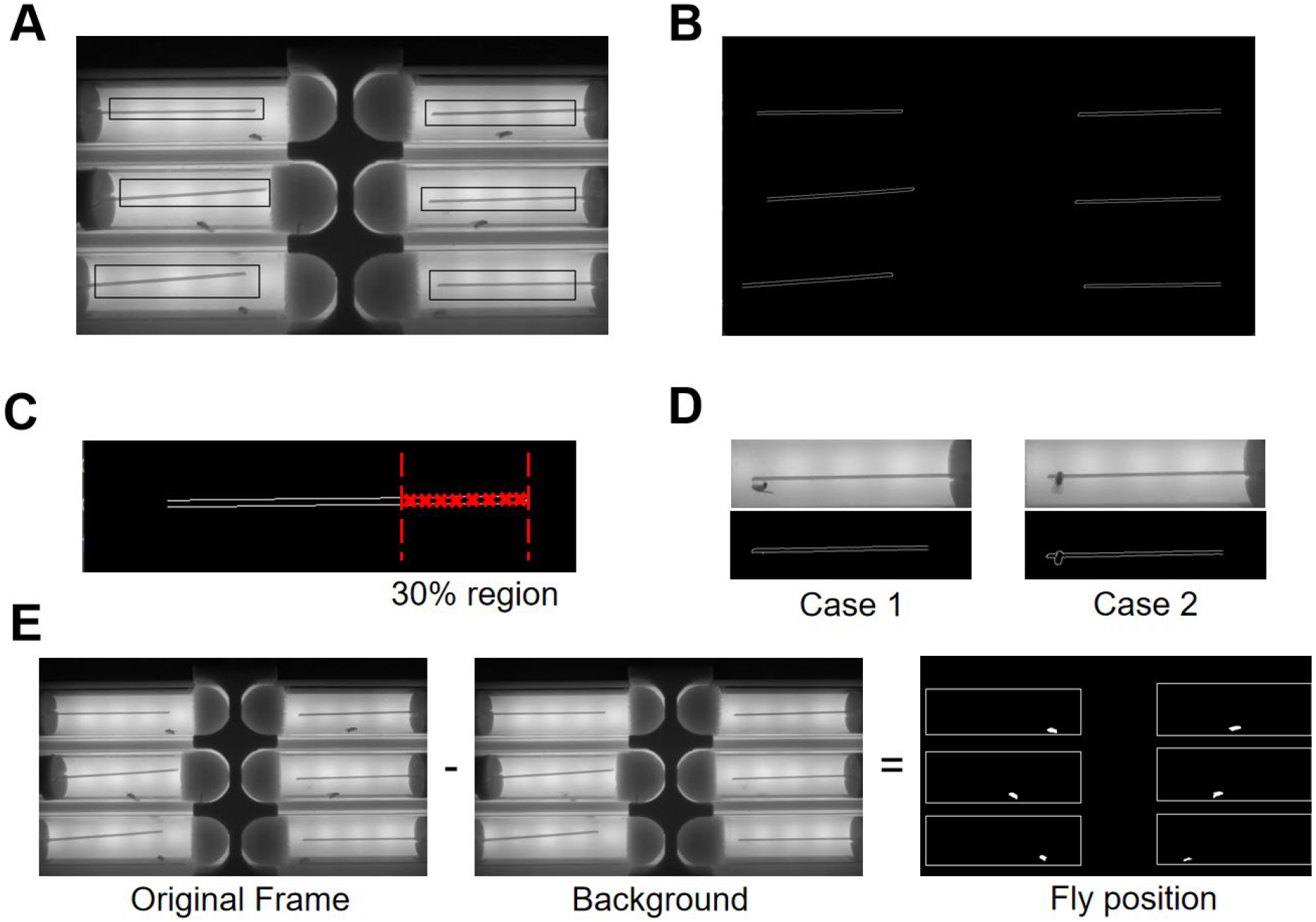
Video processing procedure for Beam and Fly Motion Detection. (**A**) Original Frame. Rectangles represent regions of interest. (**B**) The binary image containing only the edges of the beam, extracted using the Canny edge detector. (**C**) 30% region of the beam that starts from the tip is used to extract vibration information. The region is divided into 8 sections, and their centroids are calculated. The coordinates of centroids are used to extract vibration information. (**D**) Two special cases when measuring the motion signal of the beam: In Case 1, the fly stands on the beam but does not obstruct it, allowing the motion signal to be extracted without interference. In Case 2, the fly stands on the beam and obstructs it, interfering with the extraction of the motion signal. (**E**) Obtaining the binary image containing only flies using background subtraction. The background image is subtracted from the original image, and then a threshold is applied to get a binary image that only contains the fly.

We then used the background subtraction method to determine the fly’s positions. First, we extract frames from the video every 30 seconds throughout one day. By summing and averaging all extracted frames, we obtained a background image (middle image, Fig. 1E). Subtracting this background image from each video frame and applying a threshold resulted in a binary image (right image, Fig. 1E). The threshold value was chosen such that the binary image almost exclusively contained the flies. An initial threshold estimate was computed using Eqn 1 and further fine-tuned manually.

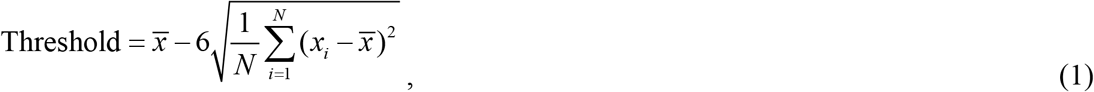

where *x* represents a set of numbers that includes all negative elements in the matrix obtained after subtracting the background image from the original video frames, and the constant 6 was an empirically determined value. To avoid interference between flies, we selected regions of interest within rectangular boxes. These regions correspond to the maneuverable area of each fly. By detecting connected components within the region, the position of each fruit fly could be determined. Once a fly’s position in the image was identified, we could determine its position on the beam by correlating the beam’s length in the image with its actual physical length.

Next, we extracted vibrations and frequencies from the beam motion signals (Fig. 2). The first step was to identify the time instances when vibrations occurred. We applied a Short-Time Fourier Transform (STFT) to the *y* coordinates of the selected centroids. Since the beam remained almost stationary most of the time, we reasoned that when vibration occurred, the energy near the natural frequency in the frequency domain should significantly increase. We located the vibration period in the motion signals by detecting energy changes. We then analyzed the vibration within this period using the Fourier Transform. Given that the vibration induced by flies lasted for a short duration, which resulted in low-frequency resolution when applying the Fourier Transform, we included some stationary signals before and after the vibration to obtain a clearer, more accurate peak. Finally, we fitted the frequency peaks using a Single-Degree-of-Freedom (SDOF) system model, yielding a more precise vibration frequency estimation, as described in Supplementary Materials, Section II.

**Figure 2.**
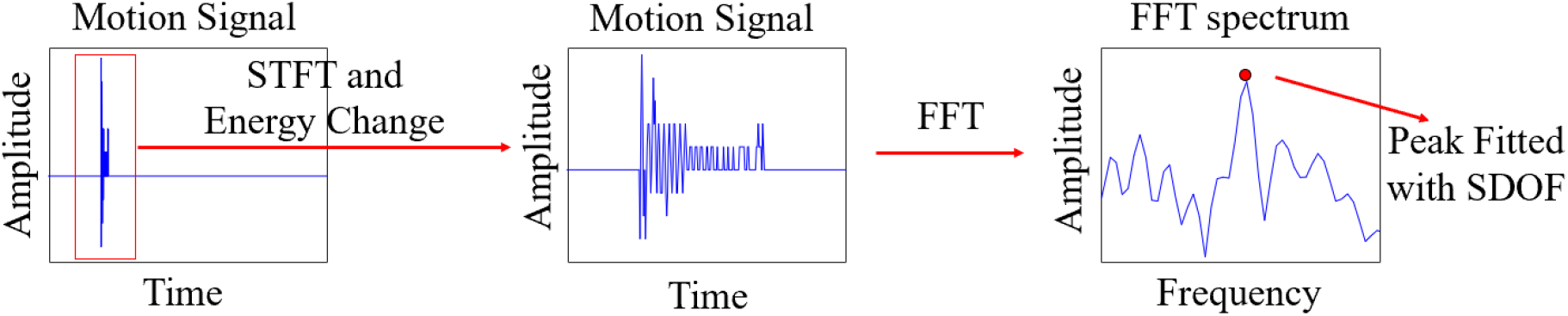
Overview of obtaining frequencies from the motion signal. The Short-Time Fourier Transform (STFT) is applied to the motion signals. The vibration events are located by checking energy change. For each vibration event, the frequencies are obtained with Fourier Transform and Single-Degree-of-Freedom (SDOF) system fitting.

### Body mass calculation

After extracting frequencies, we determined the fly’s body mass using formulas derived from the Euler-Bernoulli beam theory. The vibration system was considered to consist of an Euler-Bernoulli beam and a fruit fly, as illustrated in Fig. 3. The beam has a rectangular cross-section, fixed at one end and free at the other, with length *L*, width *b*, and thickness *h*. To simplify the model, we assumed the beam material to be isotropic, homogeneous, and vibrating linearly. The basic principle of body mass calculation is that when a fly lands on the beam and causes it to vibrate, the natural frequency of the beam decreases. The fly’s body mass can then be calculated from the reduced frequency.

**Figure 3.**
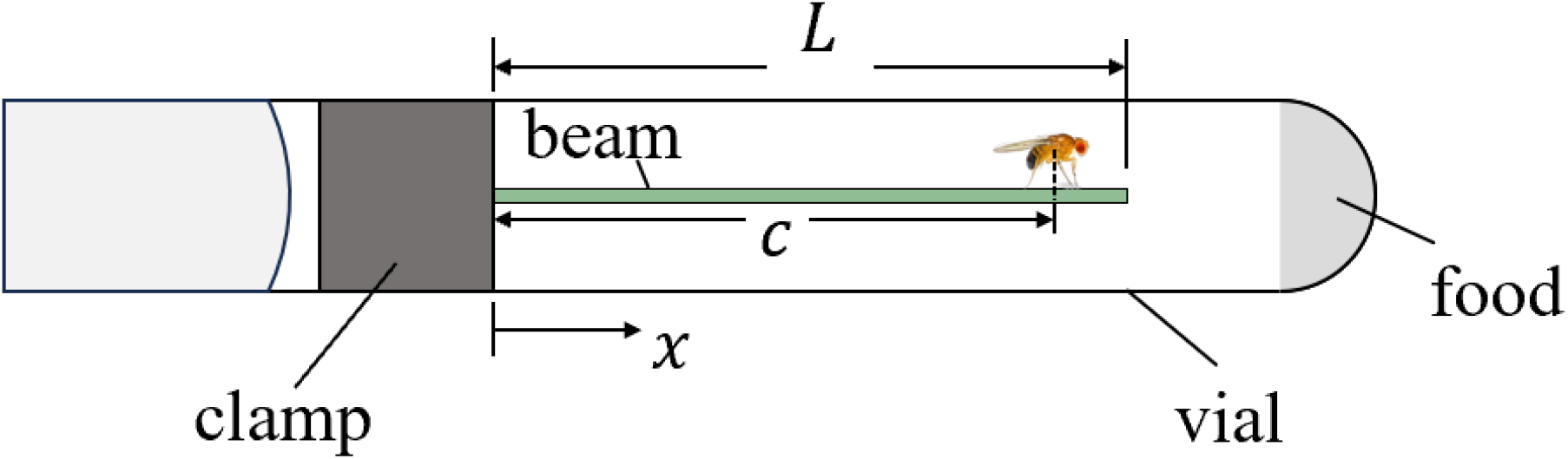
Schematic diagram of the beam-fly vibration system. The beam has a rectangular cross-section, fixed at one end and free at the other. The beam has a length *L*, width *b*, and thickness *h*. The fly can be considered a static point mass located at the position *x* = *a* on the beam during vibration.

The formula for body mass calculation is:

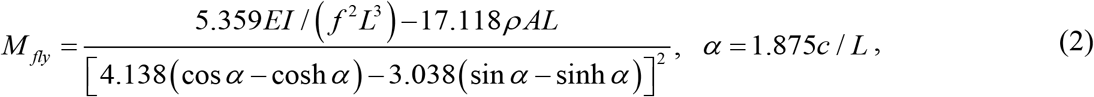

where *f* is the extracted fundamental frequency, *ρ* is the density of the beam, *A* is the cross-section area, *E* is Young’s modulus, 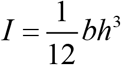 is the moment of inertia, and *c* is the position of the fly.

## RESULTS AND DISCUSSION

We used our method to monitor the body mass of wild-type *Drosophila* over a period of 14 days. An example of the tracking can be seen in Supplementary Video 1. Fig. 4 presents the estimated group-level mean trajectory of body mass over these 14 days, computed from irregularly sampled time series data collected from individual flies. To generate the trajectory, each data point was temporally aligned and interpolated onto a common time grid, followed by pointwise averaging across 5 flies.

**Figure 4.**
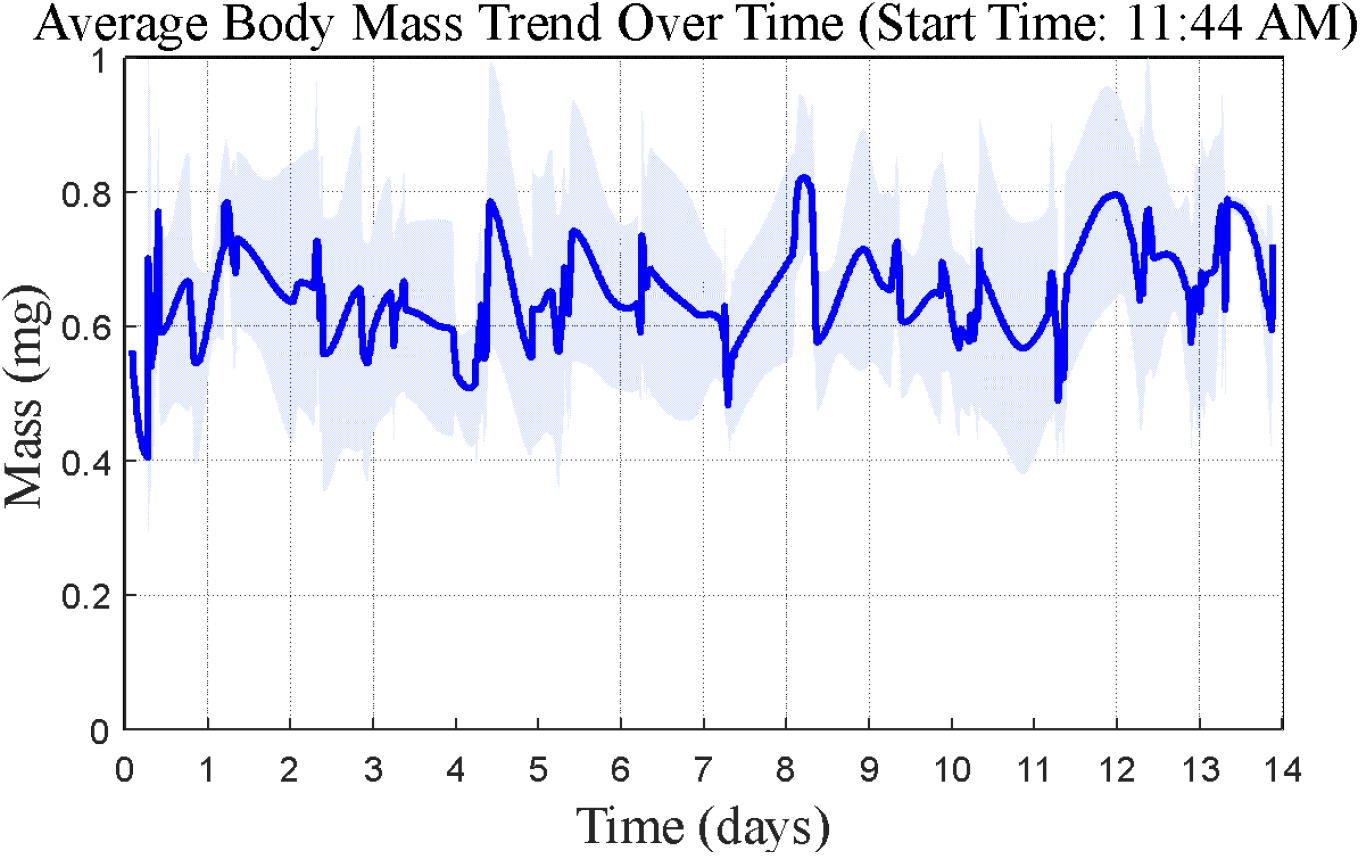
Mass monitoring experiment results. The blue curve is the average body mass trend, and the shaded area indicates the standard deviation at each time point.

The results show that the system successfully captured the fly’s body mass, with an average value of approximately 0.651 mg for wild-type individuals. This value is consistent with previously reported measurements obtained using conventional methods, thereby validating the accuracy of our system. The body mass of individual flies fluctuated within a day but remained stable overall throughout the monitoring period. This observation aligns with the established understanding that, under normal conditions, *Drosophila* body mass remains relatively constant after emergence from the pupal stage.

The system’s performance can be evaluated based on the Euler-Bernoulli beam theory. The system achieved an approximate resolution of 0.5 Hz/mg, which is sufficient for detecting small changes in fruit fly body mass. To further enhance the resolution, the system can be optimized by carefully selecting the cantilever beam’s material, refining the beam’s dimensions, and employing higher-quality cameras. Eqn 2 can be used as a guideline for optimizing cantilever beam design for improved performance.

While the system demonstrated high accuracy, several potential sources of error may have affected the measurement. The primary limitation arose from the brief duration of vibrations induced by the flies. Since these vibrations only lasted a short period, the frequency resolution obtained from the Fourier Transform was relatively low. Although we employed the SDOF system fitting to improve frequency estimation, achieving precise mass calculations remained challenging. Another potential source of error was the position of the fly on the beam. In some cases, the fly may partially obstruct the beam, interfering with the edge detection process. While our algorithm accounted for this by selecting unobstructed points for frequency extraction, minor inaccuracies may still occur. A potential solution would be to utilize the MEMS technology to fabricate beams with integrated miniature sensors, such as micro strain gauges or piezoelectric sensors, to directly measure vibrations with high sensitivity and spatial resolution.

The proposed system is highly scalable and adaptable for a variety of applications beyond monitoring fruit fly body mass. While the current design is optimized for small organisms like *Drosophila*, the underlying principles can be extended to larger animals by scaling the dimensions of the cantilever beams. For small-scale beams, the cost of conventional vibration sensors is prohibitively high. However, for larger animals, common vibration sensors such as piezoelectric sensors can replace cameras to achieve better accuracy in frequency measurement. The system can support a wide range of intriguing research projects, including studies on the relationships between genetics, disease, and body mass, as well as investigations into the interplay between genetics, diet, and body weight. Furthermore, the system’s ability to capture high-resolution images allows for the integration of behavioral analysis with mass monitoring. This system would enable researchers to explore how body mass relates to specific behaviors, such as feeding, mating, or locomotor activity in flies. By combining physiological and behavioral data, the system could facilitate more comprehensive biological research, providing deeper insights into the complex interactions among these factors.

In addition to measuring body mass, future iterations of the system could incorporate other *Drosophila* behavior monitoring technologies to further expand its functionality. For instance, additional cameras could be integrated to capture detailed behavioral patterns; acoustic sensors could be employed to record sounds generated during fly activities; and sleep deprivation modules could be introduced to facilitate studies on rest-related behaviors. These enhancements would broaden the system’s applications in both biological and behavioral research.

## Conclusion

In conclusion, we developed a cost-effective system for long-term monitoring of fruit fly body mass. The system leverages vibration analysis and image processing techniques to achieve accurate, non-invasive measurements of the body mass of individual flies over extended periods. Our experimental results confirmed the system’s ability to measure body mass, providing valuable insights into fly physiology. With its scalability and potential for integration with additional sensing technologies, this system represents a versatile platform for a wide range of biological research applications.

## ACKNOWLEDGEMENTS

We thank Jingjing Yan, Brissa Castillo, Fiona Gugala, Jennifer Dong, and Kelsey Mainard for their assistance with fruit fly husbandry. We thank Xin Chen for the comments on the manuscripts.

## DATA AND RESOURCE AVAILABILITY

The data and code used in this study are available from the corresponding authors upon request.

