## Supplementary Materials for "A Long-Term *Drosophila* Body Mass Measurement System"

**Section I. The Schematic diagram of the measurement device.**

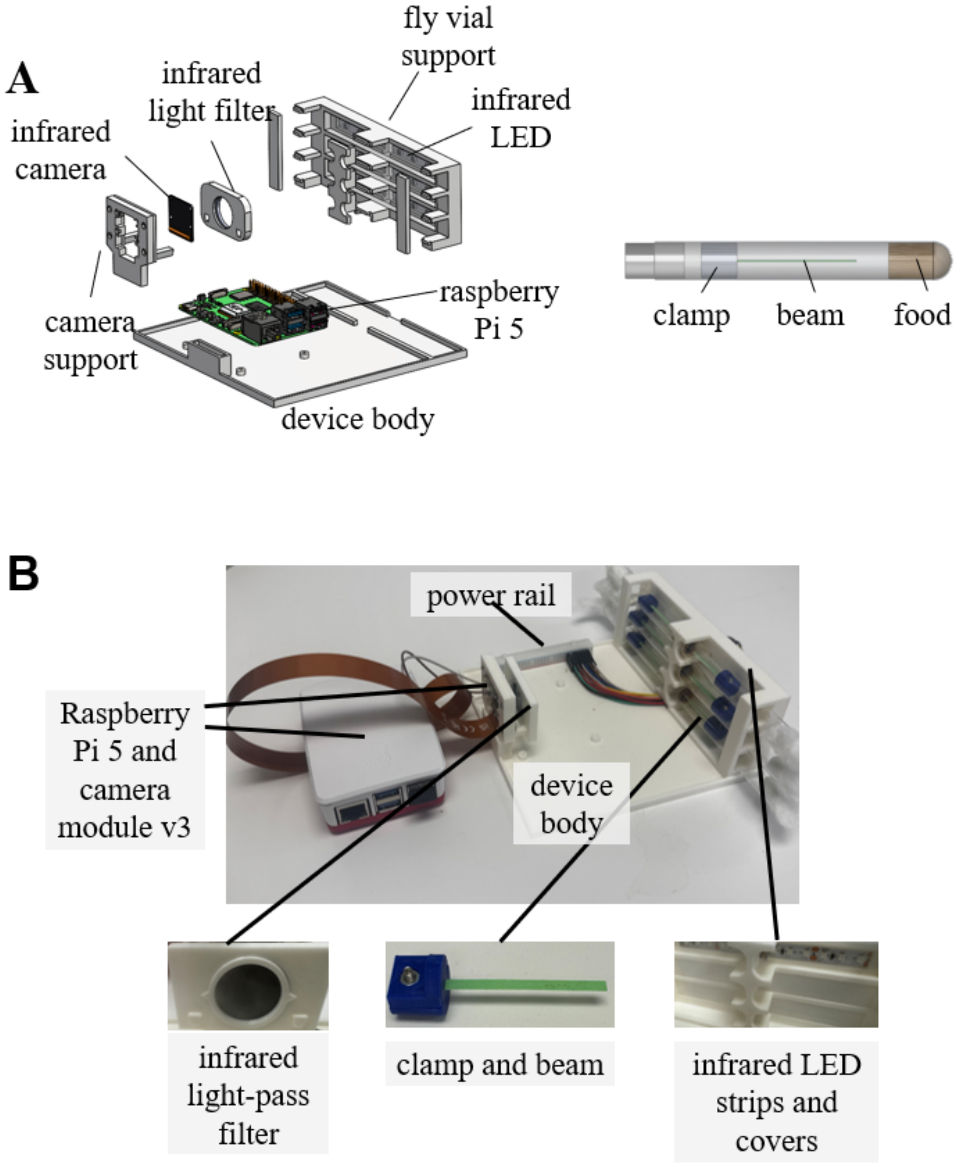


**Fig. S1** Monitoring device design. (**A**) The device is constructed using a Raspberry Pi 5 connected to a Raspberry Pi Camera Module v3 (Raspberry Pi, United Kingdom) with an 830 nm infrared light-pass filter (Thorlabs, RGL830, USA) mounted in front of the camera. The enclosure is 3D-printed using PLA filament (Ultimaker PLA 2.85 mm, Netherlands) on an Ultimaker S3 (Ultimaker, Netherlands). Each vial slot contains a 940 nm infrared LED strip (360DigitalSignage, SMD3528, China) positioned at the base, with all strips wired to the Raspberry Pi's GPIO pins via a power rail (REXQualis, 4 Breadboard Kit, China) to supply power. White acrylic covers (McMaster Carr, 8505K747-8505K817, USA) are installed above the LED strips to ensure even illumination. These covers are precision-cut using a Glowforge Plus laser cutter (Glowforge, USA). While the Raspberry Pi's 5V output is below the rated voltage of the LED strips, the emitted brightness is sufficient, eliminating the need for an additional power source. The cantilever beams are made from green polyester plastic film (McMaster Carr, 9513K15, USA), cut into shape using the Glowforge Plus laser cutter, and fixed with 3D-printed PLA clamps. The beam dimensions are $3\times10^{-2}\text{m}$ by $2.97\times10^{-3}\text{m}$ by ك $7.62\times10^{-5}\text{m}$. (**B**) Physical picture of the device.

**Section II. Fitting of peak frequency in the Fourier Transform spectrum with a Single-Degree-of-Freedom system.**

This section describes how the frequency of a peak was estimated by fitting the Fourier Transform spectrum using a Single-Degree-of-Freedom vibration system. For the peak to be fitted, the maximum amplitude is first identified, along with its half-power bandwidth (see Fig. S2), which is the frequency range where the amplitude equals the peak maximum divided by $\sqrt{2}$.

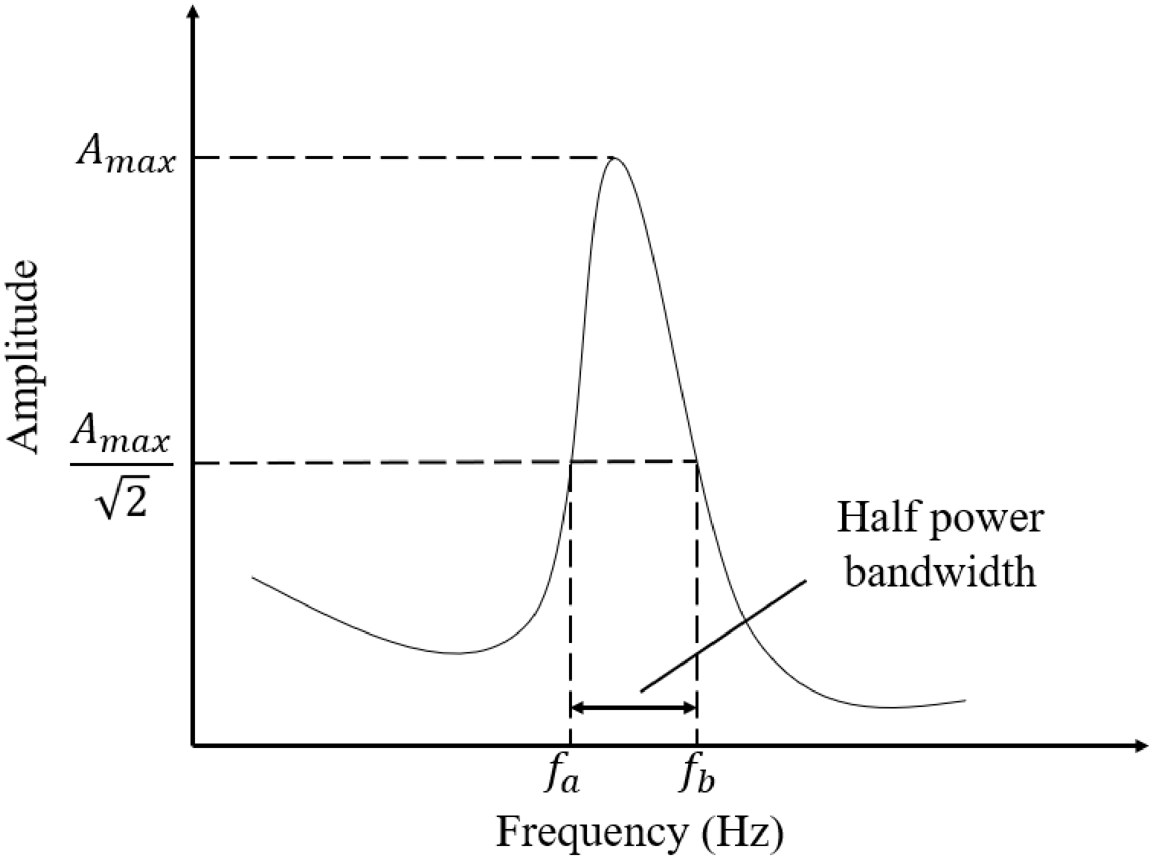


**Fig. S2** Half-power bandwidth in Single-Degree-of-Freedom fitting.

The spectrum within this range is then interpolated, and the resulting frequency–amplitude pairs are substituted into the following equation:

$$\begin{bmatrix} 1 & \frac{1}{(2\pi f_1)^2} & \frac{1}{(2\pi f_1)^4} \\ 1 & \frac{1}{(2\pi f_2)^2} & \frac{1}{(2\pi f_2)^4} \\ \vdots & \vdots & \vdots \\ 1 & \frac{1}{(2\pi f_n)^2} & \frac{1}{(2\pi f_n)^4} \end{bmatrix}_{n\times 3} \begin{Bmatrix} a_1 \\ a_2 \\ a_3 \end{Bmatrix}_{3\times 1} = \begin{Bmatrix} \frac{1}{(A_1)^2} \\ \frac{1}{(A_2)^2} \\ \vdots \\ \frac{1}{(A_n)^2} \end{Bmatrix}_{n\times 1}, \tag{S1}$$

where $a_1, a_2, a_3$ are the parameters to be estimated, $n$ is the number of points, $f_i$ is the frequency, and $A_i$ is the corresponding amplitude. The $a_1, a_2, a_3$ can be solved with the least square method and are substituted into the following equation to get the estimated frequency of the peak.

$$f_e = \frac{1}{2\pi}\left(\frac{a_0}{a_4}\right)^{\frac{1}{4}}. \tag{S2}$$

**Section III. Supplementary Video 1. An example of tracking wild-type *Drosophila*.**
